# Multiple Effects of Artificial Lighting at Night on Male Glow-worms’ Mate Searching Behaviour

**DOI:** 10.1101/2025.02.18.638807

**Authors:** Estelle M. Moubarak, A. Sofia David Fernandes, Alan J. A. Stewart, Jeremy E. Niven

## Abstract

Artificial lighting at night (ALAN) has been identified as a driver of insect decline, disrupting their ecology, behaviour and physiology. Recent studies have begun to identify the mechanisms by which ALAN affects nocturnal insects but focus primarily on moths. In glow-worms (Lampyris noctiluca), population survival relies on males detecting and reaching the glow emitted by bioluminescent females at night. Despite evidence suggesting many deleterious effects of ALAN on their mating success, little is known about the behavioural mechanisms underlying those effects. Using a translational approach, we assessed males’ ability to detect and approach females, both in nocturnal conditions and under ALAN showing that males land near a female before walking towards it. Thus, males navigate through dense vegetation to find a mate. Males walking on a trackball were highly reliable in adjusting their course to rapid left to right shifts of a green LEDs mimicking females’ glow, reorienting in less than 1 second. Under ALAN, males’ reliability to detect the dummy female was significantly hindered, along with their speed, stamina and orientation. This indicates ALAN has multiple simultaneous impacts on the ability of males to reach females, impairing their mating with potentially severe consequences on glow-worm’s population survival.

## Introduction

Artificial lighting at night (ALAN) is an increasingly widespread phenomenon, with deleterious effects on many aspects of nocturnal and crepuscular animals’ ecology including birds, mammals and insects (for reviews see ^1,2,3–5)^. The effects of ALAN are not restricted to urban areas; since the 1990s, light pollution has doubled near protected areas and in high biodiversity hotspots ^6–9^ and is predicted to keep rising. Over the last decade, several studies have established ALAN as an overlooked but key factor contributing to insect declines. Sources of ALAN can be local (*e.g.* street or car lighting) or at broader scales (*e.g.* skyglow) with the potential to affect ecosystems at local and broader scales. ALAN has been shown to disrupt aspects of insect ecology and survival (for reviews see ^3–5)^. Examples of the consequences of exposure to ALAN include the gathering of insects (primarily) at streetlights, where one third of them die before morning from exhaustion or predation (*e.g.*^10,11^), as well as delayed and less profitable foraging (*e.g.*^12,13^) and reduced mating and reproductive success (e.g. ^14–17)^.

Some studies have examined the behavioural, developmental and physiological effects of exposure to ALAN (*e.g.*^18–20)^. Insights from behaviour can reveal the mechanistic underpinnings of ecological-level observations and have led to the interpretation of long-established observations (e.g. ^20^). Most behavioural studies tend to focus on a single aspect of insects’ behaviour but a few do report multiple effects of ALAN on behaviour. In dung beetles, for example, light pollution affects the overall walking direction of individuals, through a combination of increased tortuosity and reduced orientation accuracy^21^. Even in these cases, quantification of ALAN effects on insects is often at a broader scale due to the challenges of the field setting and the spatial scales on which behaviours may occur. One means of circumventing these limitations is to simulate ALAN in laboratory settings, but these often restrict insects to small spatial scales within arenas (e.g. ^22^). Within laboratory settings, trackballs permit insects to move over large distances while allowing detailed quantification of movement at small scales (e.g. ^23^) but these have not previously been used to explore the effects of ALAN.

Males of the common glow-worm, *Lampyris noctiluca*, use phototaxis to detect and fly towards the yellowish green (546 nm) bioluminescent glow emitted by stationary females perched on plant stems^24^. This reliance on a bioluminescent display as a mating strategy makes insects such as glow-worms and fireflies and particularly vulnerable to the presence of ALAN because it can directly affect conspecific recognition, leading to a decline in mating success^17,18,25–29^ and, likely through concomitant reductions in reproductive success, is implicated in population declines (*e.g.* ^30,31^). While these studies link ALAN with reduced mating success, the majority use counts of males (and females in the case of fireflies) attracted by LED traps under different lighting conditions. Few studies, to our knowledge, investigate the specific behavioural effects of ALAN to determine the mechanisms underpinning this reduction in mating success. Female glow-worms exposed to ALAN reduce the duration of their glowing period^32,33^ whilst simulated ALAN (sALAN) in a Y-maze decreases the proportion of males initiating walking towards a dummy female through altered mate recognition and negative phototaxis^22^. In both cases, ALAN disrupts behaviours that lead to mating. Because female glow-worms are stationary, and glow for only a brief period, males must rapidly detect the glow and navigate towards it through a complex environment of dense vegetation, during which they might frequently lose sight of the female, to achieve mating. As such, characterising changes in males’ behaviour under ALAN is essential to understand how it affects glow-worms’ mating success and, consequently, population survival.

To determine the effect of ALAN on male searching behaviour for females, we adopted a translational approach. This involved recording and analysing males’ landing and approach to an LED (acting as a dummy female) in their natural habitat, as well as observing males reaching a dummy female in a behavioural arena. We combined these observations with fine-scale tracking of tethered males’ movements towards a dummy female whilst walking on a trackball in open-loop in the absence and presence of sALAN. We characterised males’ detection of and ability to reach dummy females in these simulated environments, quantifying aspects of their search strategy and the impact of sALAN upon it. We show that male glow-worms are highly effective in detecting and orientating towards the female glow but that sALAN interferes with their ability to track the glow, orientate to it and walk towards it. This shows that ALAN has the potential to disrupt multiple aspects of a critical behaviour associated with reproduction in an insect.

## RESULTS

### Male glow-worms land in the vicinity of a dummy female before walking towards it

To test whether males primarily fly, walk or combine both to reach a female, we recorded males landing in their natural habitat (Figure 1A,B; see methods). Of 170 males attracted to the trap, 126 (74.1%) reached the dummy female (Supplemental Figure 1), whilst 44 (25.9%) remained in or left the near vicinity of the trap (Figure 1C; One-sample proportion test; X^2^ =38.5, df=1, p<0.001***). Ninety four percent of males landed near the trap (Figure 1D; 46.5% >10 cm, 31.2% 5-10 cm, 16.5% <5 cm). However, just 10 males landed within 5 cm of the dummy female; seven on the rim of the trap (4%) and three entering the trap directly (1.8%) (Figure 1D). Of the 126 males that reached the trap, 117 combined walking with short flights to approach the dummy female (93%), including 111 that climbed up the side of the trap (95%), whereas only six males flew from the ground to the rim of the trap (5%). Overall, males landed on average 9.22 [5.68-10.93] cm (median [Q1-Q3]) from the LED (Figure 1E) and walked most of the distance to reach the dummy female. Thus, our field observations suggest that only 1.8% of males reach females using flight exclusively, most males combining walking with flying.

**Figure 1.**
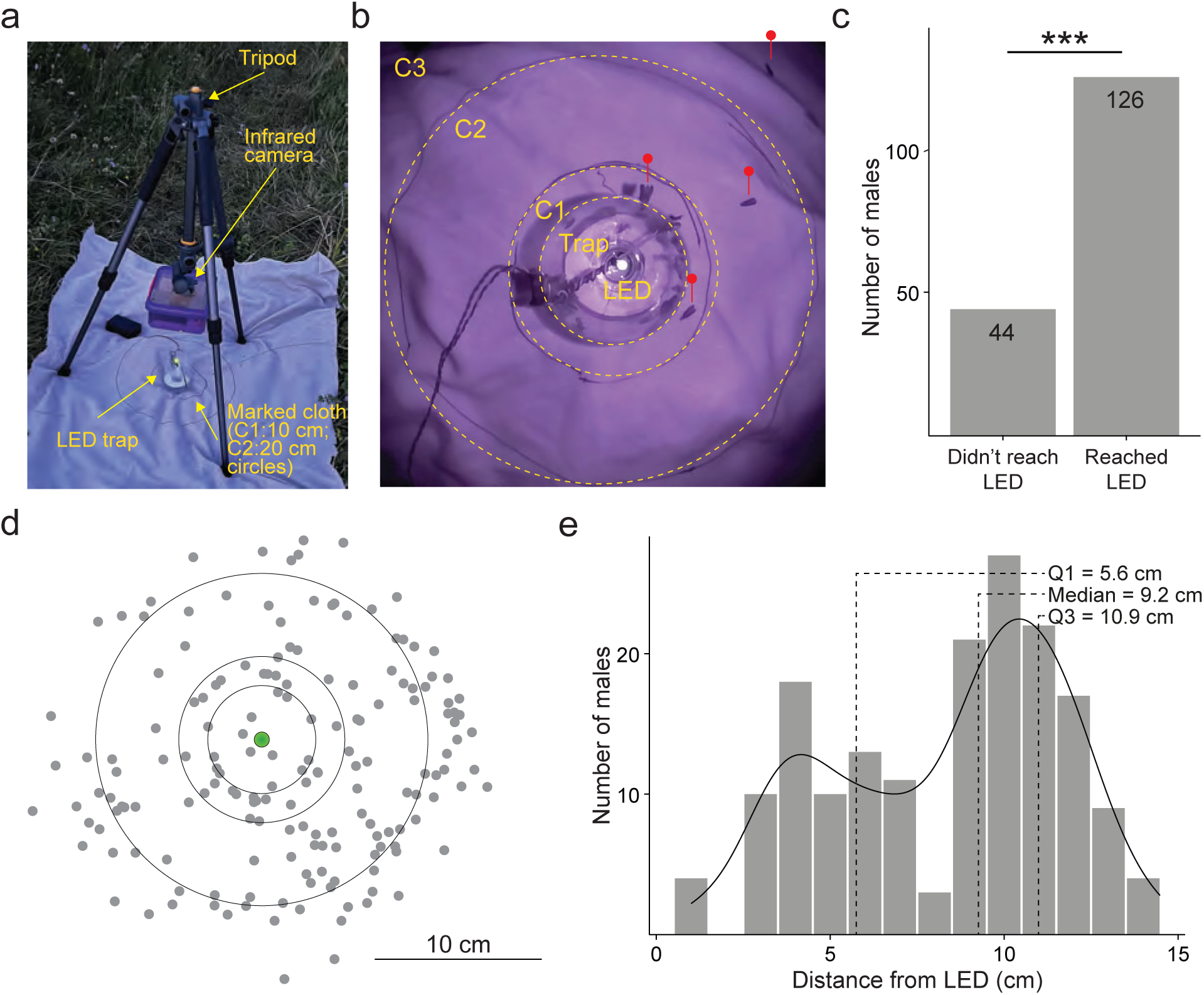
Males fly and walk to reach females in the wild. **a.** Photograph of the infrared camera setup placed in the field. The camera was connected to a Raspberry pi single-board computer was placed in a sealed box mounted face-down on a tripod above a single green LED attached to a trap. A beige cloth with circles delimiting 10 (C1) and 20 (C2) cm diameter around the LED was placed beneath the trap (see Methods). **b.** A still taken from a video recording showing the LED, the trap and the measured areas (C1, C2 and C3; see Methods). Red markers indicate the location of 4 male glow-worms in the vicinity of the trap and one on the rim of the trap. **c.** Count of males from videos that reached the LED or did not. Significance (***, p<0.001) is that of a one-sample proportion test between the two groups. **d.** Spatial locations of male landing positions on and around the LED. Grey dots represent males landing on the cloth. **e.** Distribution of males’ landing distance from the dummy female. Dotted lines indicate from left to right the 25th percentile (Q1), median, and 75th percentile (Q3). Black line is a smoothed representation of the count data.

### Freely moving male glow-worms in an arena combine flight and walking to reach glowing females

Of the 22 males placed individually in the arena (Figure 2A; see Methods), 10 leaped from the platform, whilst nine flew and three climbed down from the platform (Figure 2B). Two males flew directly to the pillar that held the dummy female, the remining 20 walking at least part of the way (Figure 2B). Five males reached the base of the pillar and 17 climbed to the dummy female. Of those males reaching the pillar, the majority did so in less than a minute (Time to pillar; 48±10 s; mean ± standard deviation) (Figure 2C). Of the males that reached the dummy female, two landed directly upon it, four males climbed the pillar directly, the remaining males (N=11) walking around the pillar before climbing to the dummy female (Time to LED; 116±24 s) (Figure 2C). Overall, there was no significant difference in the distance males flew or walked (Wilcoxon rank sum test; Flew *versus* Walked; W=102.5, p=0.7, N=22). Males that flew initially reached the pillar base significantly faster than those that leaped or climbed down from the platform (Wilcoxon rank sum test; Flew *versus* Leaped-Climb-down/Walked; W=7, p<0.001***, N=22, Figure 2D). Males walked at 1±0.3 cms^-1^ (mean ±standard deviation) while they flew at 7±4 cms^-1^. Thus, although flight allows male glow-worms to travel faster, most males walk to reach females at least over short distances and as observed in the field.

**Figure 2.**
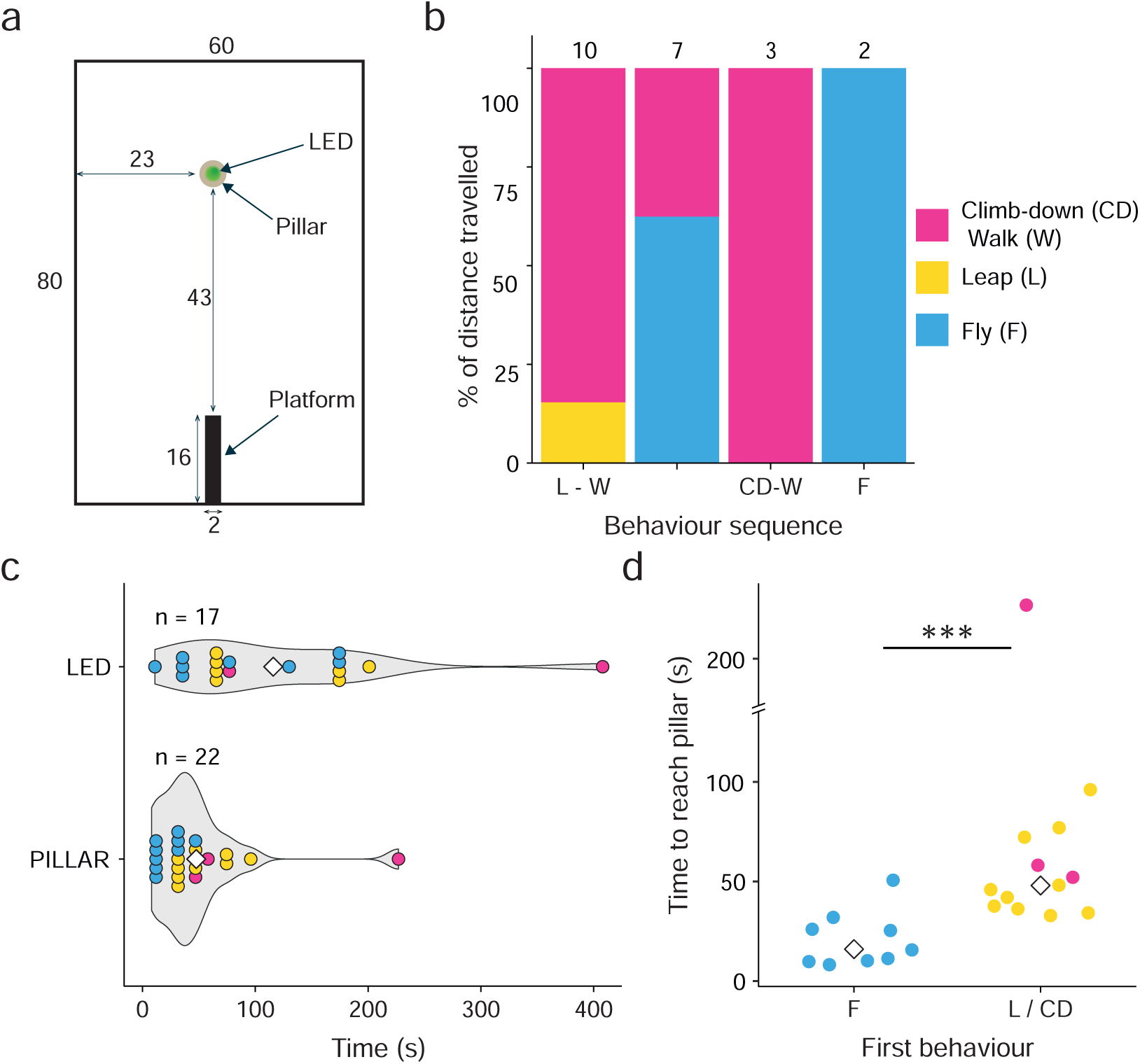
Freely moving males in an arena fly and walk to reach females. **a.** Schematic of the arena from above. All measurements given in centimetres. **b.** The average percentage of the distance to the pillar travelled by male glow-worms during each behaviour within the four observed behavioural sequences displayed as a stacked bar plot. Numbers at the top indicate the total number of males in each category. **c.** Violin plots showing males’ latency to reach either the pillar or the green LED. Dot colours correspond to those in **b.** White diamonds indicate the mean of each group. **d.** Males that climbed down or leaped from the platform took significantly longer to reach the pillar than those that flew. Dot colours correspond to those in **b.** White diamonds represent the mean of each group. Significance (***, p<0.001) is that of a Wilcoxon rank sum test comparing the time to reach the pillar. The single outlier does not affect the significance.

### Walking males reliably track the dummy female at frequencies up to 1 Hz in the dark

Males walking on the trackball (Figure 3A,B) typically responded to the presence of the dummy female by continuously attempting to walk towards it throughout the trial through both rotational and translational movement (Figure 3C,D). When the dummy female switched to the opposite side after six seconds, the male glow-worms likewise switched their direction of travel. To determine how quickly males adjust to a change in the dummy female position, they were presented with three LED switching sequences in the dark, each consisting of changes in LED position at frequencies ranging from 0.03 to 2 Hz (Supplemental Figure 2). We quantified the ability of male glow-worms to turn in the same direction as the dummy female on each trial within the overall switching sequence. Males tracked the dummy female on 79.6% (N=30, n=750), 79.1% (N=41, n=861) and 83.5% (N=17, n=85) of trials in sequence ***i*** (Figure 4A), ***ii*** (Figure 4B) and ***iii*** (Figure 4C), respectively. Males successfully tracked the dummy female location in >80% of trials at frequencies between 0.03 and 0.5 Hz (n=1128), and 71% of trials at 1 Hz (n=568). The probability of males’ tracking the dummy female was significantly dependent upon the switching frequency, higher frequencies reducing the probability of tracking (Generalised linear mixed model with Bernoulli distribution; N=110, n=2246; Figure 4G; Table 1). This marked decrease in male glow-worms responding at the highest frequency suggests an upper limit on their ability to track the female glow.

**Figure 3.**
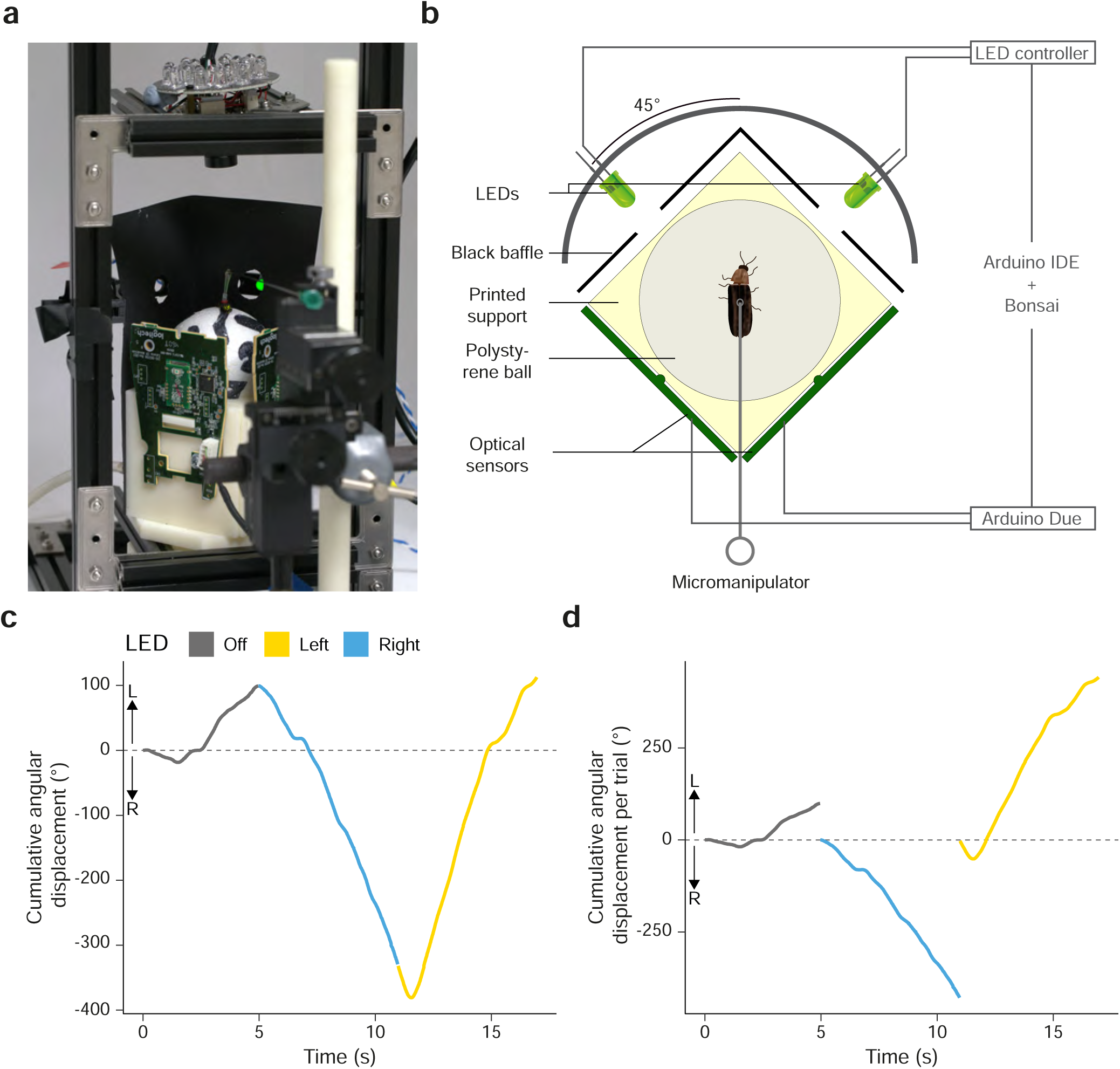
Male glow-worms walk toward a dummy female (green LED) in the dark. **a.** Photograph of the trackball setup. A male glow-worm is tethered via the micromanipulator and walking on the polystyrene ball. Optical mouse sensors can be seen attached to two sides of the ball housing. An infrared camera films the glow-worm from above. One of the green LEDs used to produce the switching patterns is visible through windows cut in the black baffle. **b.** A schematic of the trackball and its connections seen from above (see Methods). **c.** An example trace showing the path of a male glow-worm on the trackball experiencing a switch in LED position from right to left; each trial is 6 s. Male rotation to the left is shown as positive cumulative angular displacement, whereas rotation to the right is shown as negative. Colour coding of the path indicates the position of the lit LED relative to the male over time. **d.** As in **c.** but with the cumulative angular displacement per trial to facilitate the visualisation of the male orientation when the lit LED changes position.

**Figure 4.**
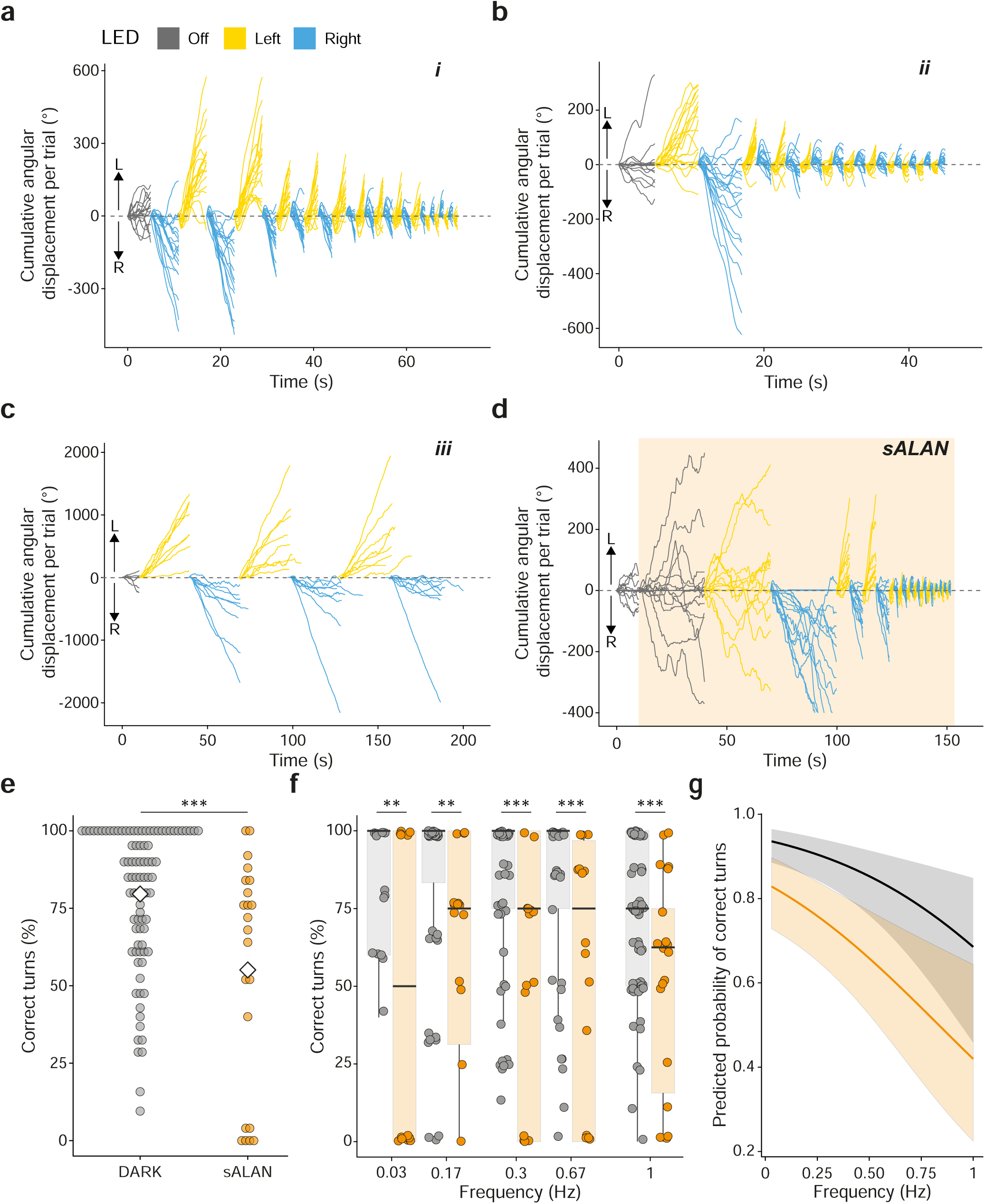
The frequency dependency of males’ ability to successfully rotate towards females in the dark nd when exposed to sALAN. **a.** Overlayed cumulative angular displacement by trial of males walking on the ackball exposed to LED switching pattern i in the dark (N=18 of 30 shown). **b.** As in **a.** but for LED switching attern ii (N=19 of 41 shown). **c.** As in **a.** but for LED switching pattern iii (N=8 of 17 shown). **d.** As in **a.** but nder sALAN (N=16 of 22 shown). **e.** The percentage rotations to the correct side for each male in the dark rey dots) or with sALAN (orange dots). White diamonds represent the mean. The significance (***, p<0.001) that of a G-test. **f.** Boxplots plots showing the proportion of rotations to the correct side per frequency for all ials in the dark and under sALAN. Boxplots represent the median, 25th percentile (Q1) and 75th percentile Q3). Dots correspond to the individual percentage rotations to the correct side for each male in the dark (grey) r under sALAN (orange). Significance (**, p<0.01; ***, p<0.001) is that of a G-test comparing the percentages etween dark and sALAN at each frequency, adjusted for multiple comparisons (FDR). **g.** Generalised linear ixed model of the Bernoulli family predicting the probability of males turning towards the green LED per equency. Solid black and orange lines indicate model prediction in the dark and in sALAN respectively. The orresponding grey and orange regions show the 95% confidence interval. See Table 1 for significance levels the model.

**Table 1.** Bernoulli Generalised linear mixed model with logit link and fit by maximum likelihood formula : Correct turns ∼Frequency + LightCondition + (1|GW)

|  | Estimates | Std. Error | Z | P |
| --- | --- | --- | --- | --- |
| **Intercept** | 2.7 | 0.033 | 8.3 | <0.001*** |
| **Frequency** | -1.1 | 0.228 | -4.8 | <0.001*** |
| **LightCondition** | -1.9 | 0.551 | -3.6 | <0.001*** |

GW = individual glow-worm. Estimate, Standard error, Z-value and significance is given for fixed effects. \*\*\*, $p < 0.001$ .

### Males’ tracking is hindered by sALAN at every frequency

Exposure to sALAN significantly reduced the percentage of males tracking the dummy female to 55.1% (N=22, n=550) (Figure 4D), a 24% reduction in comparison to males tracking in the dark at the same frequencies (79%, N=88, n=1516) (Figure 4E; G-test of independence; All trials (dark) *versus* All trials (sALAN); G=110, df=2, N=110, p<0.001***). This reduction in males tracking the dummy female under sALAN was significant at all frequencies tested (Figure 4F; Pairwise G-test, FDR-adjusted for multiple comparisons, Supplemental Table 1). As observed in the dark, the percentage of males tracking the dummy female under sALAN was significantly dependent upon the switching frequency (Generalised linear mixed model with Bernoulli distribution; N=110, n=2246; Figure 4G; Table 1). Thus, the presence of sALAN affects males’ ability to detect and approach females in a similar manner at every tested frequency.

### sALAN increases males’ latency to turn towards the dummy female

We measured the latency of male turns towards the dummy female following a switch (Supplemental Figure 3A). In the dark, males turn towards the dummy female in under a second (Figure 5A; 0.59 [0.46-0.73] s; median [95% confidence interval]). Exposure to sALAN, increased the males’ latency (Figure 5A; 0.62 [0.44-0.82] s). The effect of sALAN on males’ latency was significantly different in low frequency (0.03-0.5 Hz; N=110, n=631, W=23604, p<0.001, Figure 5B) but not in high frequency trials (Figure 5B). Indeed, males’ latency to respond was frequency independent in the dark (Generalised linear mixed model with Gaussian distribution; N=110, n=1150; Table 2; Figure 5C). Conversely, under sALAN males’ latency decreased with increasing frequency (N=110, n=1150; Table 2; Figure 5C) possibly due to adaptation to the ambient white light or as a consequence of only the most efficient males responding to the highest frequency trials.

**Figure 5.**
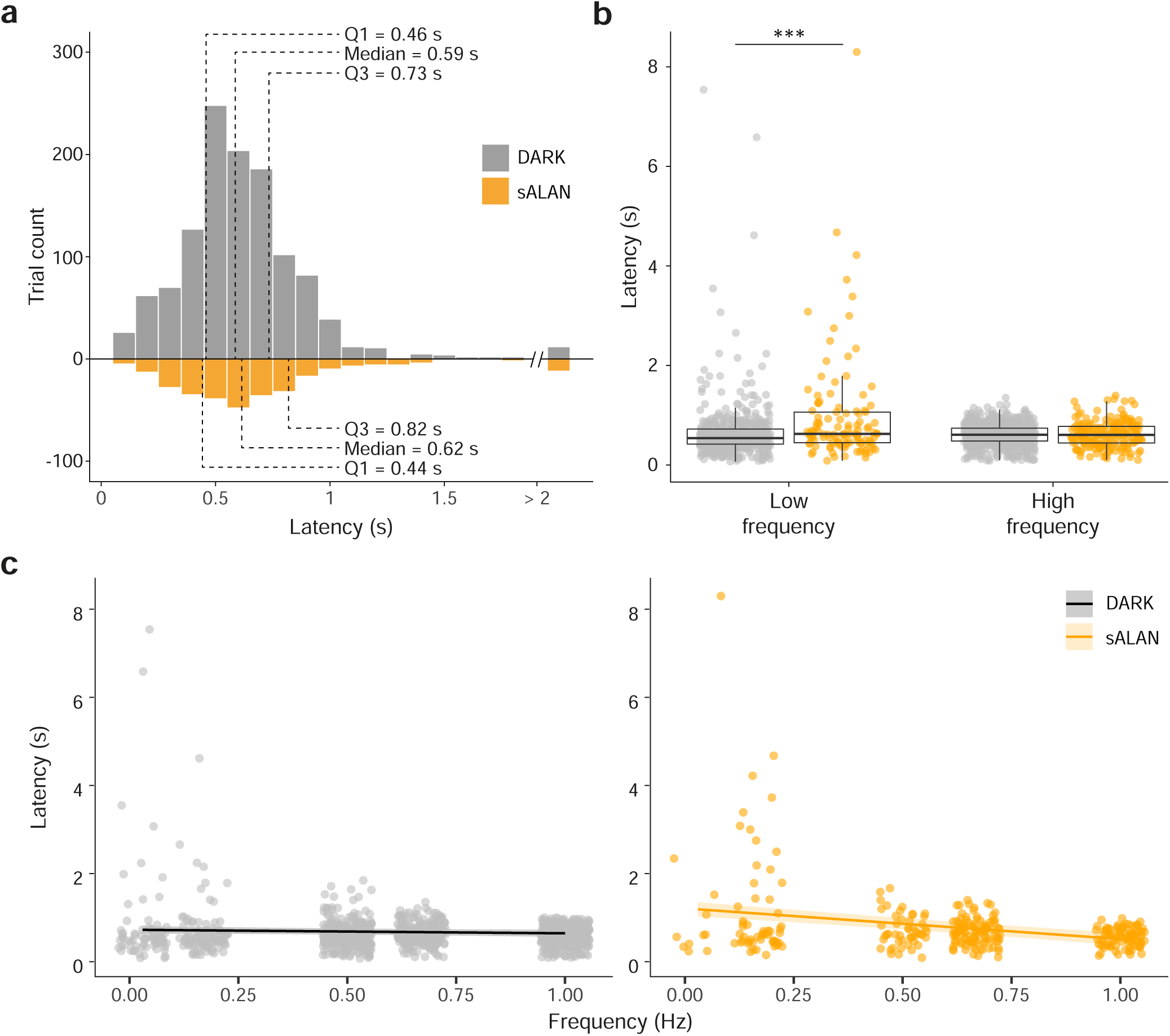
The latency with which male glow-worms’ paths change during a switch in LED position is pid even in the presence of sALAN. **a.** Distribution of males’ latency to initiate a rotation towards the green ED in the dark (grey bars) and in sALAN (orange bars). Dotted lines indicate from left to right the 25th percen-e (Q1), median, and 75th percentile (Q3). **b.** Boxplots showing the latency to initiate a rotation between males the dark and males in sALAN during lower frequency (0.03-0.5 Hz; left) or higher frequency (0.67-1 Hz; right) witches. Boxplots represents the median, 25th percentile (Q1) and 75th percentile (Q3). Dots correspond to e individual latency measurement for each male on each successful trial in the dark (grey) and in sALAN range). Significance (***, p<0.001) is that of a Wilcoxon sum rank test comparing latency between both oups. **c.** Linear mixed model of the latency to turn towards the dummy female at each tested frequency in the ark and in sALAN. Solid black and orange lines indicate model prediction in the dark and in sALAN respective-. The corresponding grey and orange regions around the line corresponds to the 95% confidence interval. ots indicate individual recorded latencies in the dark (grey) and sALAN (orange). See Table 2 for significance vels of the model in the dark and sALAN.

**Table 2.** Gaussian Linear mixed model fit by restricted maximum likelihood of formula : Latency Frequency + LightCondition + Frequency:LightCondition + (1|GW)

|  | Estimates | Std. Error | Z | P |
| --- | --- | --- | --- | --- |
| **Intercept** | 0.72 | 0.044 | 16.5 | <0.001*** |
| **Frequency** | -0.08 | 0.048 | -1.68 | 0.093 |
| **LightCondition** | 0.48 | 0.094 | 5.17 | <0.001*** |
| **Frequency:LightCondition** | -0.61 | 0.097 | -6.35 | <0.001*** |

GW = individual glow-worm. Estimate, Standard error, Z-value and significance is given for fixed effects. \*, $p < 0.05$ ; \*\*, $p < 0.01$ ; \*\*\*, $p < 0.001$ .

### Males rapidly match their orientation to the dummy female position in the dark but not under sALAN

Males on the trackball are aligned to 0 ° (facing forward) and cannot change their body angle. To head towards the dummy female, they must attempt to adjust their walk to rotate to 45 ° to the left or right. To assess the accuracy of this rotation, we calculated males’ path orientation during the initial 1 s of the first trial. In the dark, males adopted a median initial heading angle of 37 [25-55] ° (Supplemental Figure 3B; Figure 6A). In the presence of sALAN, males adopted a more acute orientation of 10 [5-11] ° (Figure 6B), significantly different from that of males in the dark (Figure 5C; Watson’s Two-Sample Test of Homogeneity; sALAN *versus* Dark; F = 0.557, N=48, n=48, p<0.001***) (Figure 6C). We also calculated males’ latency to orientate fully towards the dummy female (*i.e.* to rotate to a 45 ° angle). In the dark, males are predicted to attain a 45 ° angle in 1.2 [0.8-2.3] s (Figure 6D), whereas the presence of sALAN increases the time males are predicted to take to 3.3 [2.8-4] s; a significant difference in time taken to adopt the orientation to the dummy female (Figure 6D; Wilcoxon rank sum test; sALAN *versus* Dark; W = 37, N=48, n=48, p<0.001***). Thus, even though males exposed to sALAN can turn towards the dummy female, their ability to adopt the correct orientation is impaired.

**Figure 6.**
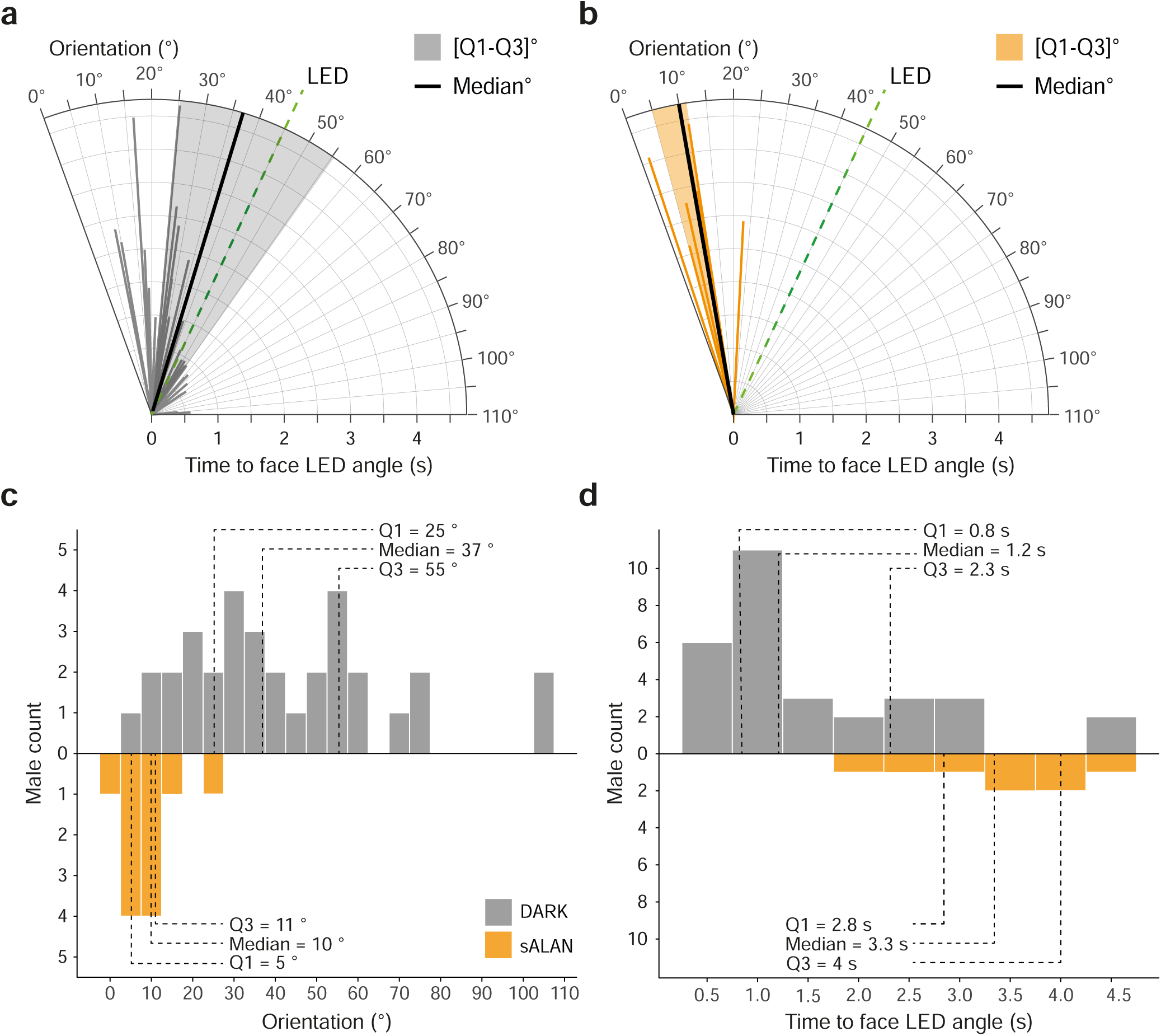
Males’ initial heading directions are more accurate in the dark than under sALAN. **a.** Polar plots howing the heading angle of each male in the dark relative to the position of the green LED. Grey lines show dividual heading angles (line angle) and time taken to face the LED (line length), the black line shows the edian and the grey sector is the interquartile range [Q1-Q3]. **b.** As **a.** but for males exposed to sALAN. **c.** istribution of males heading angles in the dark (grey bars) and in sALAN (orange bars). **d.** Distribution of tency to attain a heading direction in line with the position of the green LED in the dark (grey bars) and in ALAN (orange bars). Dotted lines indicate from left to right the 25th percentile (Q1), median, and 75th percen-e (Q3).

### Males in the dark walk faster towards dummy females than do those exposed to sALAN

After initiating a turn towards a dummy female, males’ initial angular velocity, calculated from the initial 1 s, was 33 [17-55] °/s (Figure 7A). In the presence of sALAN, the initial angular velocity was 19 [10-31] °/s (Figure 7A), 40% slower than for males in the dark, a significant difference in initial angular velocity (Wilcoxon rank sum test; sALAN *versus* Dark; W=235912, N=88, n=1470, p<0.001***). Males’ initial angular velocity in the dark was faster at every switching frequency than under sALAN except for the fastest, although this could be due to the trials being too short to accurately fit slopes (Figure 7B; Pairwise Wilcoxon sum rank test, FDR-adjusted for multiple comparisons, Supplemental Table 2). For males in the dark, the angular velocity throughout the entire trial was 31 [15-53] °/sec, which was strongly correlated with the initial angular velocity (Linear Gaussian Regression, R=0.91, N=88, n=1170, p<0.001***) (Figure 7C). Likewise, under sALAN males’ angular velocity throughout the entire trial was 18[10-28] °/s, which was strongly correlated with the initial angular velocity (Linear Gaussian Regression, R=0.89, N=15, n=300, p<0.001***) (Figure 7C). The difference in entire trial angular velocity was significant between males in the dark and under sALAN (Wilcoxon rank sum test; sALAN *versus* Dark; W=235484, N=88, n=1470, p<0.001***) and at every frequency except for the slowest (Supplemental Table 3) suggesting that given enough time, males can adapt to the presence of sALAN and increase their speed. The short 1Hz trials did not allow for accurate slope fitting.

**Figure 7.**
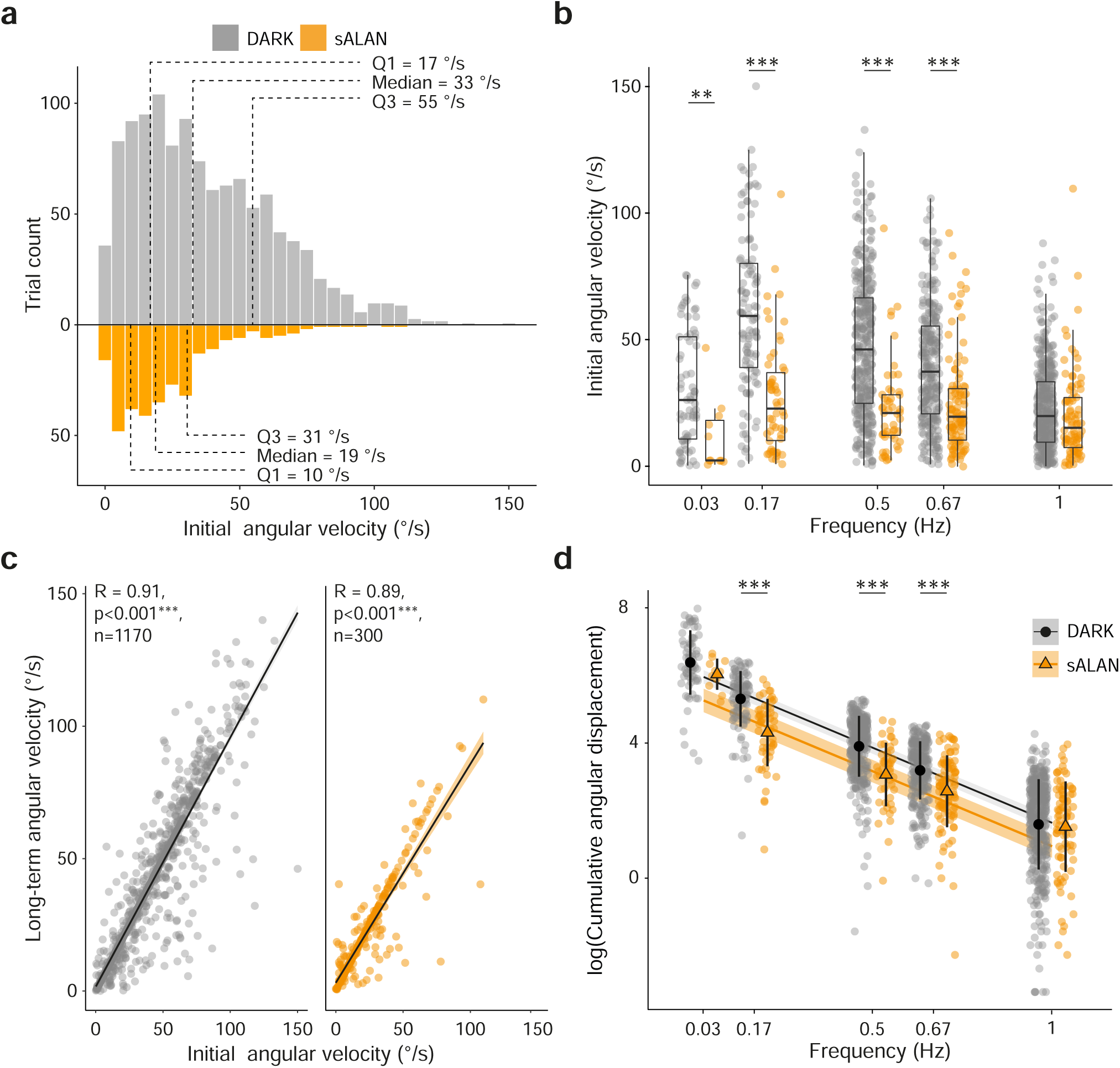
Simulated artificial lighting at night (sALAN) reduces males’ initial and long-term angular elocity in comparison to males in the dark. **a.** Distribution of male’s initial angular velocity in the dark (grey ars) and when illuminated by a white LED (sALAN). Dotted lines indicate from left to right the 25th percentile 1), median, and 75th percentile (Q3). **b.** Boxplot and individual points showing the initial angular velocity in e dark (grey dots) and under sALAN (orange dots) across a range of frequencies. The boxplot represents the edian, 25th percentile (Q1) and 75th percentile (Q3). Significance (*, p < 0.05; **, p < 0.01; ***, p < 0.001) is at of a Wilcoxon sum rank test comparing initial angular velocity between both groups at each frequency, djusted for multiple comparisons (FDR). **c.** Linear regressions showing the relationship between initial and ng-term angular velocity in the dark (grey dots, left) and under sALAN (orange dots, right). Dots show individal measures of angular velocity. **d.** Linear mixed-effects model of the relationship between the logged total ngular distance walked by a male across frequencies in the dark (grey dots) and under sALAN (orange dots). lack dots or orange triangles and vertical lines represent the mean and standard deviation, respectively. ignificance (*, p < 0.05; ***, p < 0.001) is that of a Wilcoxon sum rank test comparing the angular distance etween both groups at each frequency, adjusted for multiple comparisons (FDR).

The walking velocity of males affects the distance they cover during a trial. In the dark, males’ mean angular distance during the experiment was 52 [32-104] ° (Figure 7D), whereas in the presence of sALAN the mean angular distance walked by males was reduced to 38 [25-50] ° (Figure 7D), a significant difference in angular distance (Wilcoxon rank sum test; sALAN *versus* Dark; W=1026, N=88, p<0.05*), despite the experiments in sALAN being on average longer than the ones in the dark (sALAN experiment duration = 112 s, average dark experiment duration = 70 s; Supplemental Figure 2). The total angular distance walked by males throughout a trial depended upon the switching frequency in the dark and in sALAN (Generalised Linear Mixed model with lognormal distribution, Figure 7D, Table 3) because they have a longer period over which to walk at lower frequencies than at higher frequencies (Figure 7A-D). Males exposed to sALAN walk less than males in the dark at most tested frequencies (Figure 7D, Table 3, Supplemental Table 4). As observed for males’ velocity, this difference is not significant at the highest frequency, as both groups walk only a relatively short distance in the very limited amount of time the LED is lit. This suggests that below 1 s, even in the dark, males can initiate a rotation toward the dummy female but have insufficient time to make significant progress towards it. Interestingly, males’ also walk equivalent distances in the dark and in sALAN at the smallest frequency of 0.03 Hz, suggesting that given enough time, males might adapt to the presence of white light, as the long-term velocity also indicates. Overall, these results suggest that sALAN substantially alters males’ ability to track females, impacting not only their probability of response but also the accuracy and strength of their approach.

**Table 3.** Lognormal Linear mixed model fit by restricted maximum likelihood of formula : g(Angular distance) ∼ Frequency + LightCondition + (1|GW)

|  | Estimates | Std. Error | Z | P |
| --- | --- | --- | --- | --- |
| **Intercept** | 6.1 | 0.1 | 60 | <0.001*** |
| **Frequency** | -4.5 | 0.09 | -51 | <0.001*** |
| **LightCondition** | -0.7 | 0.2 | -4 | <0.001*** |

GW = individual glow-worm. Estimate, Standard error, T-value and significance is given for fixed effects. \*\*\*, $p < 0.001$ .

## Discussion

Our aim was to understand how male glow-worm’s reach glowing females in the dark and the impact of ALAN upon this. To achieve this, we combined observations from the field and in a behavioural arena with detailed quantitative analysis of tethered males’ movements on a trackball both in the dark and in the presence of simulated ALAN (sALAN). Our field and arena observations show that males primarily walk in the final stages of their approach to females rather than flying. On the trackball, males tracked the female glow reliably, rapidly responding to the presence of the glow and changes in its relative position. Exposure to sALAN reduced males’ ability to respond to the presence/position of the glow, but even when they did respond their paths were impaired to some extent in every aspect we quantified.

Our field videography shows that most male glow-worms (98%) reach a dummy female by walking after landing 9.2 (0-15) cm away, though this is likely an underestimate because 14% of males landed beyond our camera’s field of view. Our observations from males in a behavioural arena also show they primarily walk to reach a dummy female. In both cases (and for trackball experiments), the dummy female was a green 555nm narrowband emission spectrum closely resembling that of the female glow, which peaks at ∼546nm^34,35^. Several studies have shown that similar LEDs reliably attract males in field^25–29^ and laboratory experiments^22^. This suggests that our dummy females are suitable substitutes for female glow-worms and, consequently, that males approach females over short distances using walking. Previous anecdotal descriptions of glow-worm mating^24^ suggest that most males fly directly to the female. However, our observations suggest a more likely strategy: males detect female bioluminescence while flying, land and then walk the final distance to the female. This strategy is reminiscent of that of the potato tuber moth (*Phthorimaea operculella*); only 17% of males reach a dummy female suspended from the ceiling through flight, while 68% of males can reach one sitting on a wall by walking^36^. Male moths land 5-65 cm away from the dummy female and walk the remainder of the distance^36^. Likewise, female parasitoid flies (*Ormia ochracea*) locate male crickets using phonotaxis to deposit their larvae, landing ∼8.2 cm from a dummy cricket^37^, comparable to male glow-worm landing accuracy. This suggests that combining flight over longer distances with walking over short distances may be a common strategy in insects for approaching small stationary targets: walking permitting fine-tuning of insects’ goal-directed orientation in complex habitats, such as dense vegetation.

Our custom-made trackball system was inspired by and is similar in design to those used previously with bees^38^, ants^39^, crickets^40^, flies^41^ and locusts^42^. Despite being restricted to a forward-facing position, males tethered to our trackball walked for up to 25 minutes (EMM pers. obs.) suggesting neither trackball nor tether impaired their movement. Our approach of switching the LED from left to right at different frequencies was motivated by considering how walking males approach females: sedentary females sit on a plant stem (often in dense vegetation) to display their glow^24^, after landing males walking through the vegetation to reach the female will change position relative to the glow, requiring them to adjust their course multiple times. We assessed four aspects of males’ movements: i) turning reliability; ii) response latency; iii) orientation accuracy; and iv) angular velocity and angular distance covered. We found that male glow-worms (∼80%) consistently steer towards the dummy female location regardless of stimulus frequency. Males typically turn towards the new location of the dummy female within ∼600 ms, though some males respond with a visual detection time as short as 200 ms. Similar processing times have been found in fruit fly and locust voluntary escape or hiding behaviour from looming stimuli^43,44^, suggesting that male glow-worm responses to the position of the dummy female require similar levels of neural processing. Males’ response latency is maintained across multiple trials up to at least 1 Hz, suggesting rapid feedback from and behavioural implementation of visual information. Males achieve a median angle of 37 ° in one second and accurately orient themselves to the dummy female position of 45 ° within 1.2 s, though some do so within 500 ms. Added to a minimum visual detection time of 200 ms, this indicates a minimum processing time of 700 ms from visual signal detection to its translation into an accurately directed locomotor output. Males’ initial rotational velocity was highly correlated to their long-term velocity, suggesting that the amplitude of male’s response did not decrease over the 30 s duration of our trials. Overall, our results indicate that male glow-worms are highly reliable, fast, accurate and persistent in their search for mates.

*L. noctiluca* are capital breeders^45^, investing the energy reserves accumulated during their larval stages to fuel their reproduction as adults. In this context, energy is as a finite resource, and rapidly attaining a mate is an essential part of their reproductive fitness^46–48^. Our field observations show that >20 males can land near a single female within a short period. To ensure its mating success, a male must reach the female before its competitors. Such nonaggressive “scramble competition”^45^ is common when females are spatially dispersed and males approach them simultaneously in a limited time; both conditions are met by glow-worms’ ecology and life history. In scramble competition, the ability to detect and track a mate, as well as directed and rapid locomotion, are major determinants of mating success in several insect species, including some fireflies^49–52^. This suggests that male glow-worms’ reliability, orientation accuracy, speed and robustness is important for their mating success and, consequently, their reproductive fitness.

Our data shows that ALAN has multiple deleterious effects on glow-worm males’ ability to reach females to initiate mating. Simulated ALAN (sALAN) impairs males’ ability to initiate a turn towards the dummy female on the trackball by 24%. Even when males do turn towards the dummy female, their orientation is impaired by 27 °, their angular speed by 14 °/s and the mean angular distance they walked by 14 °. Males’ latency to initiate a turn towards the dummy female was also increased under sALAN at low frequencies. The effects of sALAN (∼5-14 Lux) in males walking on the trackball are greater than those produced by brighter sALAN (45 Lux) on males walking in a Y-maze^22^. One explanation for this is that males on the trackball are exposed to sALAN produced by LEDs with a short-wavelength narrow peak at ∼450 nm and a smaller long-wavelength broad peak around 550 nm, whereas earlier Y-maze experiments used a light source that lacks the distinct blue peak. The presence of short-wavelength light reduces the attractiveness of the female glow for male glow-worms^25,34^, and can affect abundance and feeding behaviour in other species such as moth caterpillar^53^. The sALAN to which males were exposed on the trackball is comparable in intensity and spectrum to modern white LED streetlights, which have increasingly been used to replace low- or high-pressure sodium streetlights or mercury streetlights^54–56^. Moreover, the green 555 nm LEDs that the male glow-worms were tracking are brighter (∼3 Lux) than the female glow, suggesting that the impacts of ALAN from streetlighting may be even greater than we find. Thus, our experiments suggest that modern white LED streetlighting would severely impair multiple aspects of male glow-worm mate searching and probably to a greater extent than more traditional streetlighting.

In bioluminescent insects, such as *L. noctiluca*, the impact of ALAN is likely to be particularly potent and may act as an evolutionary trap (for reviews see^3–5)^. Previous studies have shown ALAN can delay or stop females glowing thereby reducing the temporal mating window^32,33^. However, where females are exposed to ALAN but continue to glow, they can act as beacons, luring males into areas in which our results show their ability to reach females for mating will be severely impaired. Indeed, dummy female traps exposed to ALAN continue to attract males, albeit in lower numbers^22,25,27–29^. Our results explain why fewer males arrive at such traps but also raise the possibility that larger numbers of males are arriving nearby dummy females but are sufficiently impaired that they remain walking in the undergrowth without reaching the traps themselves.

Despite growing evidence of the impact of ALAN on insects (for reviews see^3–5)^, the majority of studies focus on the direct ecological consequences of ALAN rather than its effects on behavioural mechanisms, with some exceptions. In bioluminescent insects, for example, exposure to ALAN alters fireflies’ courtship flashes and mating success^57–59^. In *L. noctiluca,* it can delay or stop females glowing thereby reducing the temporal mating window^32,33^, impair males’ ability to recognise the glow and induce negative phototaxis^22^. Obtaining mechanistic insight revealing the underlying causes of ecologically observed effects has been valuable in a variety of other species. For example, recent work has identified a behavioural mechanism that can explain why nocturnal moths gather at streetlighting^20^; moths turn their dorsum towards the brightest visual hemisphere to maintain their flight altitude, producing bouts of perpendicular flights that result in circling around the light source. In dung beetles, individuals use celestial cues such as the moon and the stars to walk in a straight path to and from food sources^60^. Exposure to ALAN reduces their orientation accuracy and increases path tortuosity, potentially resulting in an increased energetic cost of travel and less efficient foraging^21^.

Studies of behavioural mechanisms can offer new insights into the impact of ALAN on insects, providing explanations of effects observed at the ecological level. Our results show that the impact of ALAN on the reduction in male glow worms arriving at females^18–22^ is likely the consequence of multiple effects of ALAN on male glow worms’ ability to reach females. In conjunction with previous research on the effect of ALAN upon females^32,33^ and males^22^, this shows that ALAN impairs numerous aspects of behaviour that precede mating in the glow worm. Such effects on mating could have serious negative consequences for the long-term survival of glow work populations.

## Methods

### Animals

Male glow-worms (*Lampyris noctiluca*; Linnaeus, 1767) were studied or collected from the Mount Caburn National Nature Reserve, East Sussex (UK).

### Field video-recordings and analysis

Videos of males landing in the field were obtained using a Raspberry Pi Zero 2 W (Raspberry Pi Ltd, Cambridge, UK) connected to a night vision infrared camera module with a wide-angle lens (160 °) (Pimoroni Ltd, Sheffield, UK), a power bank (PB-HOUMI-1W; Elefull, Shenzhen, China) and a screen (4inch-HDMI-LCD; Waveshare, Shenzhen, China). All electronics were placed in a sealed Perspex box (L20*W15*H8) along with silica gel desiccant beads to keep electrical components dry. The camera box was mounted on a tripod (Alta Pro 2+, Vanguard, Dorset, UK), with the lens sitting approximately 36cm above the floor (Figure 1A). We used a custom-made LED trap consisting of a green 555 nm LED (Supplemental Figure 1; SSL-LX5093PGD, Lumex Inc., Carol Stream, IL, USA) mounted above a funnel trap tapering from 8 to 2 cm^22^. The LED was powered by three 1.5 V batteries and faced upward at approximately 12 cm from the ground to allow it to sit beneath the camera (24 cm between camera and LED). A beige cotton sheet was placed below the trap to permit detection of the males landing on the ground. The sheet was marked with two concentric circles of 10 and 20 cm diameter around the LED (Figure 1A, B) to permit estimation of landing distance.

We recorded five 1-hour long videos (30fps) on five separate nights, all starting at approximately 21:50 and ending at 22:50. Videos were opened using Fiji/ImageJ^61^ and the “VideoImportExport” plugin (https://sites.imagej.net/VideoImportExport/) for offline analysis. A total of 170 males were observed across all videos. Males’ landing location was marked and classified according to the area they first landed in (directly in the trap, on the trap rim, or on the ground in either C1, C2, or C3; Figure 1A, B). Males were then followed until they either entered the trap or exited the field of view. We also scored whether males that landed on the ground flew into the trap or climbed up the side of the trap to reach the dummy female (LED).

### Capture, maintenance and identification of males

Males were captured using custom-made traps (as described above) but with the LED at approximately 20 cm from the ground. After capture, males were brought into the laboratory where they were marked with acrylic paint dots on the elytra for subsequent individual identification and placed in clear Perspex boxes (L22*W13*H14 or L17*W11*H5) in a dark room with a controlled day/night cycle of 10:14 hour dark:light (dark from 13:00 to 23:00). Illumination was produced with cool white LED strips (ATOM LED lighting, Telford, UK) plugged on programmable timer switches. The plastic containers contained holes in the lids to permit air circulation and were lined with wet paper towels to maintain a high humidity. A small shelter of black cardboard allowed the glow-worms to hide from the light. For all trackball experiments, glow-worms were kept in these conditions for at least 48 hours prior to any experiment.

### Arena video-recordings and behavioural classification

To assess males’ preferred means of travelling towards a female, we constructed a rectangular arena (L80*W60*H60 cm) containing a platform (L16*W2) raised 4 or 9 cm above the ground and a single green 555 nm LED (as described above) raised on a pillar 3 cm above the ground and 43 cm from the platform to mimic female glow (Figure 2A). At the start of each experiment, an individual male was placed on the platform and the LED turned on immediately. All experiments were performed in dim light conditions and videos were recorded under infrared light using a USB infrared camera (ELP-USBFHD05MT-RL36-U, Shenzhen Ailipu Technology Co., China). Recordings were stopped when the male reached the LED or if more than 15 minutes had elapsed since the male initiated a movement. Data from both platform heights were grouped because no discernible difference in the glow-worms’ behaviour could be detected.

We analysed males’ mode of travel by assessing the behaviours they performed to dismount from the platform and then to travel towards the dummy female. Behaviours were classified into: “Leap”, males that opened their wings upon dismounting the platform but covered less than 30% of the distance between the platform and LED (13 cm); “Fly”, males that opened their wings upon dismounting the platform and flew more than 30% of the distance; or, “Climb-down”, males that climbed down the platform to the arena floor. A further behavioural classification “Walk” described males that walked with wings closed between the platform and LED. The final location of males was classified as “LED” if they reached the pillar and climbed up to the LED or “Pillar” if they reached the vicinity of the pillar (2 cm radius circle around the LED, Figure 2A) but did not climb. Of the 28 males tested, those that remained on the platform (N=5) or fell from it (N=1) were excluded from further analysis.

The videos of individual trials were analysed offline using the Fiji/ImageJ software^61^. For the males retained for analysis (N=22); we calculated approach time from the initiation of the first behaviour to the moment the male reached the pillar. The time taken to climb the LED was calculated from the moment the male reached the pillar to when it contacted the LED. Time measures are given in seconds as (mean ± standard error). Distances and speed were measured as a straight line between the locations of the male at the start and end of each behaviour.

### Insect tethering

For experiments involving the trackball (see below), a small piece of clear tubing (1.02 mm ID, 1.98 mm OD, 5 mm length, Smith Medical Inc, USA) was attached on males’ elytra above the thorax using low melting point wax. This allowed us to secure a male onto a custom-made holder (a pipette tip fitting the tubing, glued to a bent syringe needle) attached to a micromanipulator^62^. This configuration prevented males from rotating when moving on the trackball (Figure 3A,B). Attachment of the tube was performed at least 24 hours before the start of an experiment.

### Trackball setup

A tethered male was placed atop a light polystyrene ball (5 cm diameter) supported on an air cushion delivered through a hole in the base of a custom-made cup-shaped structure^62^. The air cushion was produced by a pump (ET 80 Air Pump, Charles Austen, UK). This allowed rotation of the ball with little friction as the glow-worm walked upon it. Ball movements were recorded by two optical mouse sensors (M500, Logitech, Switzerland) (Figure 3A,B). Each optical sensor was connected to an Arduino Due (Arduino, Italy) that recorded the x and y displacement of the ball, relaying it to a computer. A ball attached to a step motor was used to calibrate the trackball movement.

The light stimulus consisted of a pair of green 555 nm LEDs (as described above) held on a custom-made LED array placed around the trackball. These LEDs were positioned 3 cm from the trackball centre, at a 45 ° angle on the left and the right side (Figure 3A,B), and oriented to face the male placed in the centre at an intensity of 3 Lux. Simulated ALAN (sALAN) was simulated using a pair of white LEDs (Supplemental Figure 1, VAOL-5LWY4, VCC optoelectronics, San Marcos, CA, USA) placed 3 cm above the green LEDs, orientated to face the glow-worm at 5-14 Lux. The LEDs were connected to a microcontroller running Arduino IDE that permitted the stimulus intensity and duration to be set and retrieved as well as a constant current LED driver to improve LED stability and longevity (LED Zappelin^63^). A USB infrared camera (ELP-USBFHD05MT-RL36-U, Shenzhen Ailipu Technology Co., China) mounted above the trackball recorded videos of the experiments. The x and y values coming from the optical sensors, the LED status from the controller and the videos were collated and saved using Bonsaisoftware^64^.

### Trackball stimuli

Males were tethered upon the trackball in the dark for at least 2 minutes prior to being presented with a green LED stimulus (the dummy female). The males were presented with one of three LED switching patterns in the dark (Supplemental Figure 2B), or an LED switching pattern with the white LEDs switched on (sALAN, Supplemental Figure 2B). The green LED switched from left to right repeatedly, with the patterns of switching varying in trial number and frequency in the dark (Supplemental Figure 2B). In the sALAN condition, the dark adaptation period was followed by exposure to 30 seconds of white light at an intensity of 10-20 Lux. The white LEDs remained on while the green LED switching pattern was presented to males (Supplemental Figure 2B). In all conditions and switching patterns, the starting side for the green LED was randomised so that ∼50% experiments started on the left and ∼50% on the right side.

### Trackball path and variable extraction

All paths were processed and variables extracted in R 4.3.2^65^. Calibrated x and y values from the optical sensors were used to calculate the angular displacement of males in degrees and visualise their path. The overall cumulative angular displacement (Figure 3C) and the angular displacement normalised per trial (Figure 3D) were extracted. For clarity, in paths visualisations Figures 4A, B, C and D, approximately half of the males (selected randomly) are represented for each LED switching pattern, but all variable extractions and subsequent statistical analysis are performed on the entire dataset. These paths permitted extraction of several variables describing the male’s efficacy in localising and moving towards a dummy female. Initially, we classified trials as a binary ‘response’ or ‘failure to respond’, capturing whether a male responded to a green LED being turned on. A response was characterised either by an increase in activity (from a baseline of low or no activity) when the green LED was turned on or the side switched, or a clear change in a male’s orientation towards the side of the lit green LED. A failure to respond occurred if the male neither increased its activity when the green LED was turned on nor changed the direction of its path following a switch in the stimulus side. Additionally, trials were deemed a ‘failure to respond’ if the male was already walking in the direction of the green LED before a switch occurred.

For trials classified as a ‘response’, we then calculated additional parameters: males’ latency to respond to the green LED (Latency, s); their initial orientation (Orientation, °); the speed at which they rotated towards the dummy female (Initial angular velocity, °/s); the distance walked during the trial (Angular distance,°); and, the overall mean angular velocity throughout the trial (Long-term angular velocity, °/s). The latency to respond was measured in two different ways depending on the males’ behavioural state in the few seconds preceding the change in stimulus (Supplemental Figure 3A); if the male was active in the opposite direction, latency was considered to be the time at the absolute peak of the angular displacement, whereas if the male was inactive, change point modelling (cpm) was used to fit the time of change in activity using the {mcp} package^66^ (v0.3.3). To assess the accuracy of males’ rotation towards the lit green LED, we extracted from the path their initial orientation: the angle to which they rotated in the first second of responding to the stimulus on the first trial. Initial angular velocity was measured as the slope of the path during the first second after responding to the stimulus, whereas the total angular velocity was the slope throughout the trial (Supplemental Figure 3B). The angular distance walked was calculated as the sum of all distances oriented in the correct direction toward the stimulus over the entire trial (Supplemental Figure 6B).

### Statistical analysis

All statistical analyses were conducted in R 4.3.2^65^. For field video recordings, the statistical significance of the proportion of males reaching the trap was assessed using One-sample proportion test from the {stats} package^65^ (v4.3.2); against a random distribution of probability p = 0.5. In the arena analysis, comparisons of the distance walked versus flown and the time to reach the pillar in walking-leaping versus flying males were assessed using Wilcoxon Rank Sign test from the {stats} package.

For the trackball, all comparisons of ‘responded’ or ‘failed to respond’ trial outcomes were assessed by G-test from the {DescTools} package^67^ (v0.99.52). All comparisons of non-binary continuous variables (Latency, Angular velocities and Angular distances) were assessed using Wilcoxon rank test from the {stats} package. After the initial analysis of the response rate on the first two trials (Figure 3, Supplemental Figure 4), all analyses were conducted excluding the first trial because it was significantly different from subsequent trials. An exception was the initial orientation which can only be extracted on the first trial. Orientation comparisons were assessed using Watson’s Two-Sample Test of Homogeneity from the {CircStats} package^68^ (v0.2.6). Bernoulli family generalised linear mixed-effect models (GLMMs) from the {glmmTMB} package^69^ (v1.1.10) were used to compare the probability of response of males during and after a trial. Using the same package, Linear mixed-effects models following Gaussian and lognormal distribution were used to model the effects of frequency or trial number on males’ latency to rotate towards the green LED in the sALAN condition and to compare the Angular distance walked during a trial between the dark and sALAN condition across frequencies respectively. The relationships between initial and long-term velocities in the dark or sALAN conditions were assessed by Linear Gaussian Regression from the {stats} package. Where appropriate, p-values were adjusted for multiple comparisons using false discovery rate correction.

### Spectrometry

The emission spectra of the green and white LEDs (Supplemental Figure 1) were obtained using a commercial spectrophotometer (CCS100/M, Thorlabs, Newton, NJ, USA) and its dedicated OSA software. Light intensity was recorded using a commercial photometer (ALX-8809A, ATP Instrumentation, Ashby-de-la-Zouch, UK).

## Supporting information

All Supplementary material

## Acknowledgements

We thank Andre Maia Chagas and Maxime Zimmermann for their help implementing the electronics for LED stimuli on the trackball, and Kevin Brady for his help 3D printing the trackball support. We also thank Natural England and Glynde Estates for the permission to work at the Mount Caburn National Nature Reserve.

## Author contributions

Conceptualization: E.M.M., A.S.D.F., J.E.N.; Methodology: E.M.M., A.S.D.F., J.E.N.; Experimental investigation: E.M.M., A.S.D.F.; Statistical analysis – design : E.M.M., J.E.N.; Statistical analysis – implementation : E. M.M.; Data curation: E.M.M., A.S.D.F; Writing – original draft: E.M.M., J.E.N.; Writing – review & editing: E.M.M., A.S.D.F., A.J.A.S., J.E.N.; Visualisation: E.M.M., J.E.N.; Project administration: E.M.M., J.E.N.; Funding acquisition: A.J.A.S., J.E.N.

## Data availability

The data and analysis script are available in the supplementary information.

## Supplementary information

Supplemental information, including three figures and four tables, is available online (INSERT JOURNAL LINK). All data files associated with this study are available online (Temporary link: https://datadryad.org/stash/share/wYp8xXDQZWNOixLjs1t06jzohZcbPyZIJiNnsyFEapY). Additional information about the trackball setup is available online at https://github.com/Sussex-Neuroscience/NL-glow-worm/tree/main.

