## Supplementary material for "Multiple Effects of Artificial Lighting at Night on Male Glow-worms’ Mate Searching Behaviour": All Supplementary material

### Supplementary Tables & Figures

**Supplementary Table 1. Pairwise G-test of independence comparing the count of rotations to the correct side in the dark vs. in sALAN at each frequency**

| Frequency (Hz) | G | FDR-adjusted P |
|---|---|---|
| **0.03** | 9.8 | <0.01** |
| **0.17** | 10.1 | <0.01** |
| **0.5** | 31 | <0.001*** |
| **0.67** | 32 | <0.001*** |
| **1** | 30 | <0.001*** |

G-value and significance are given per frequency. **, p<0.01; ***, p<0.001. FDR, false discovery rate.

**Supplementary Table 2. Pairwise Wilcoxon Sum Rank test comparing the initial angular velocity in the dark vs. in sALAN at each frequency**

| Frequency (Hz) | W | FDR-adjusted P |
|---|---|---|
| **0.03** | 597 | <0.01** |
| **0.17** | 4665 | <0.001*** |
| **0.5** | 12846 | <0.001*** |
| **0.67** | 18524 | <0.001*** |
| **1** | 19637 | 0.06 |

W-value and significance are given per frequency. **, p<0.01; ***, p<0.001. FDR, false discovery rate.

**Supplementary Table 3. Pairwise Wilcoxon Sum Rank test comparing the angular velocity throughout an entire trial in the dark vs. in sALAN at each frequency**

| Frequency (Hz) | W | FDR-adjusted P |
|---|---|---|
| **0.03** | 489 | 0.196 |
| **0.17** | 4499 | <0.001*** |
| **0.5** | 12693 | <0.001*** |
| **0.67** | 18398 | <0.001*** |
| **1** | 19637 | 0.06 |

W-value and significance are given per frequency. ***, p<0.001. FDR, false discovery rate.

**Supplementary Table 4. Pairwise Wilcoxon Sum Rank test comparing the angular distance walked by males in the dark vs. in sALAN at each frequency**

| Frequency (Hz) | W | FDR-adjusted P |
|---|---|---|
| **0.03** | 511 | 0.103 |
| **0.17** | 4624 | <0.001*** |
| **0.5** | 12735 | <0.001*** |
| **0.67** | 18073 | <0.001*** |
| **1** | 18645 | 0.675 |

W-value and significance are given per frequency. ***, p<0.001. FDR, false discovery rate.

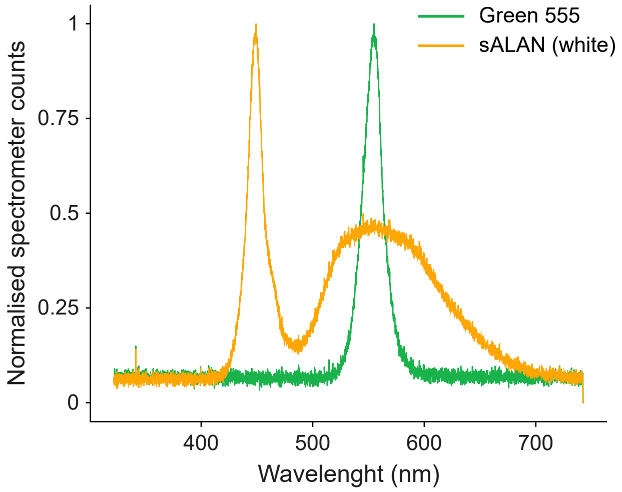


**Supplementary Figure 1. The emission spectra of the green LED used as a dummy female and the white LED used to create sALAN.** Normalised emission spectra of the green LED (green line) used to mimic the glow of female glow-worms and white LED (orange line) used to simulate artificial lighting at night.

**a**

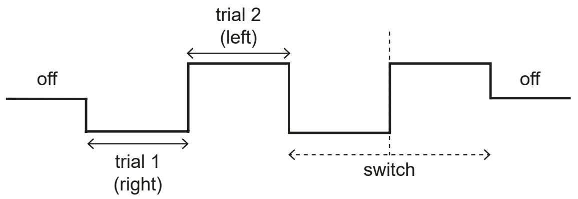


**b**

| Switching pattern | | | Stimulus duration (s) | Number of trials |
|---|---|---|---|---|
| DARK | *i* | (Hz) 0.17 0.33 0.5 1<br>(s) 6 3 2 1 | 66 | 26 |
| | *ii* | (Hz) 0.17 0.5 0.67 1<br>(s) 6 2 1.5 1 | 40 | 22 |
| | *iii* | (Hz) 0.03<br>(s) 30 | 180 | 6 |
| *sALAN* | | (Hz) 0.03 0.17 0.5 0.67 1<br>(s) 30 6 2 1.5 1 | 112 | 26 |

**Supplementary Figure 2. Protocols used to control the green LEDs during trackball experiments. a.** Graphical representation of a stimulation pattern showing our nomenclature. A trial represents a single LED position (left or right, plain arrows), while a switch indicates a change in LED position (dotted line) and the two trials surrounding it (dotted arrow). **b.** Table describing the LED switching patterns used in the experiments. The left-hand column indicates whether the pattern of simulation was used in the dark (three switching patterns i-iii) or during simulated artificial lighting at night (sALAN). The central column shows the switching pattern. Initially the green LEDs are 'off' and then to 'on' either on the left or right of the male glow-worm. Numbers indicate the duration that the green LED remained on during a trial in seconds (s) and the corresponding frequency (Hz). The grey overlay represents an overhead white light LEDs being switched on (sALAN). The right-hand columns show the total duration of a single pattern of switching and the number of trials that glow worm-males experienced.

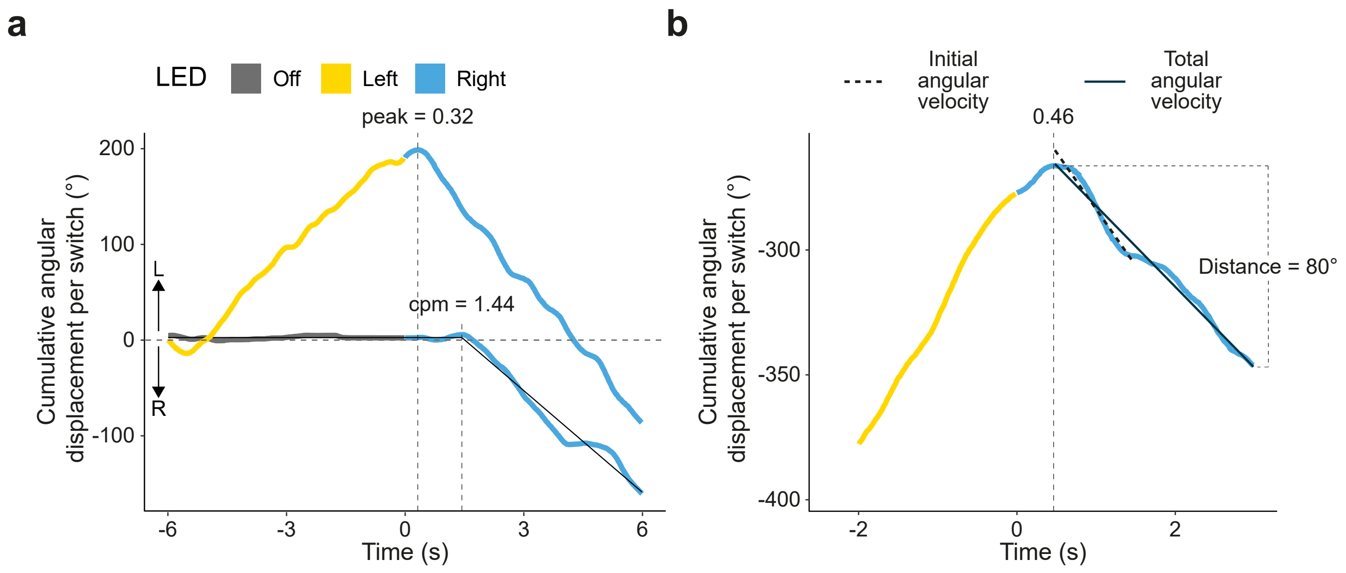


**Supplementary Figure 3. Detection and measurement of trackball parameters. a.** Examples of two cumulative paths with dashed vertical lines showing the latency calculated from peak detection (peak) or change-point modelling (cpm) (see Methods). **b.** An example of a cumulative path illustrating the total distance walked towards the LED (dashed bracket), the initial angular velocity (dashed regression line), the total angular velocity (plain regression line) and the latency (dashed vertical line) (see Methods).
